# Cortical Connectivity In A Macaque Model Of Congenital Blindness

**DOI:** 10.1101/188888

**Authors:** Loïc Magrou, Pascal Barone, Nikola T. Markov, Herbert Killackey, Pascale Giroud, Michel Berland, Kenneth Knoblauch, Colette Dehay, Henry Kennedy

**Affiliations:** Univ Lyon, Université Claude Bernard Lyon 1, Inserm, Stem Cell and Brain Research Institute U1208, 69500 Bron, France; Université De Toulouse Paul Sabatier, 31062 Toulouse, France; Centre De Recherche Cerveau & Cognition, CNRS, UMR 5549, 31059 Toulouse, France; Princeton Neuroscience Institute and Department of Psychology, Princeton University, Princeton, 08544, USA; Department of Neurobiology and Behavior, University of California, Irvine, Irvine, California 92717, USA

## Abstract

Brain-mapping of the congenitally blind human reveals extensive plasticity(*1*). The visual cortex of the blind has been observed to support higher cognitive functions including language and numerical processing(*2, 3*). This functional shift is hypothesized to reflect a metamodal cortical function, where computations are defined by the local network. In the case of developmental deafferentation, local circuits are considered to implement higher cognitive functions by accommodating diverse long-distance inputs(*4–7*). However, the extent to which visual deprivation triggers a reorganization of the large-scale network in the cortex is still controversial(*8*). Here we show that early prenatal ablation of the retina, an experimental model of anophthalmia in macaque, leads to a major reduction of area V1 and the creation of a default extrastriate cortex (DEC)(*9, 10*). Anophthalmic and normal macaques received retrograde tracer injections in DEC, as well as areas V2 and V4 post-natally. This revealed a six-fold expansion of the spatial extent of local connectivity in the DEC and a surprisingly high location of the DEC derived from a computational model of the cortical hierarchy(*11*). In the anophthalmic the set of areas projecting to the DEC, area V2 and V4 does not differ from that of normal adult controls, but there is a highly significant increase in the relative cumulative weight of the ventral stream areas input to the early visual areas. These findings show that although occupying the territory that would have become primary visual cortex the DEC exhibits features of a higher order area, thus reflecting a combination of intrinsic and extrinsic factors on cortical specification. Understanding the interaction of these contributing factors will shed light on cortical plasticity during primate development and the neurobiology of blindness.

## Main Text

Early visual cortex deafferentation via bilateral removal of the eyes at early stages of prenatal development in the macaque provides a non-human primate (NHP) model of anophthalmia. In NHP anophthalmics, there is an in-depth modification of the development of the visual system accompanied by characteristic sulci malformations; cortex destined to become striate cortex (area V1) reverts to a default phenotype (Default Extrastriate Cortex DEC)(*9, 10,12*). The three anophthalmics used in this study each showed important shifts in the border of striate cortex (area V1) leading to an important reduction in its dimensions (**Fig. 1**). In these animals instead of the typical border between areas V1 and V2 one can detect a large region of interceding cortex where the stria of Genari is absent and its cytoarchitecture can be broadly defined as extrastriate. This stretch of cortex exhibits small islands of striate cortex and the hybrid expression of histochemical phenotypes of striate and extrastriate cortex both during *in utero* development and postnatally(*12, 13*). The cortex between area V2 and the reduced striate cortex area V1 corresponds to the DEC (lower panels of **Fig. 1A,B**). The proportion of area V1 with respect to the total cortex is considerably reduced in anophthalmic brains compared to the normal (**Fig. 1C**). This contrasts with the proportions of total visual cortex (including DEC) with respect to neocortex, which is similar in anophthalmics and normals, therefore coherent with deafferentation causing a border shift rather than merely a shrinkage of area V1. Hence the DEC plus the remaining area V1 in the anophthalmic matches the extent of area V1 in the normal animal(*14*).

**Fig. 1.**
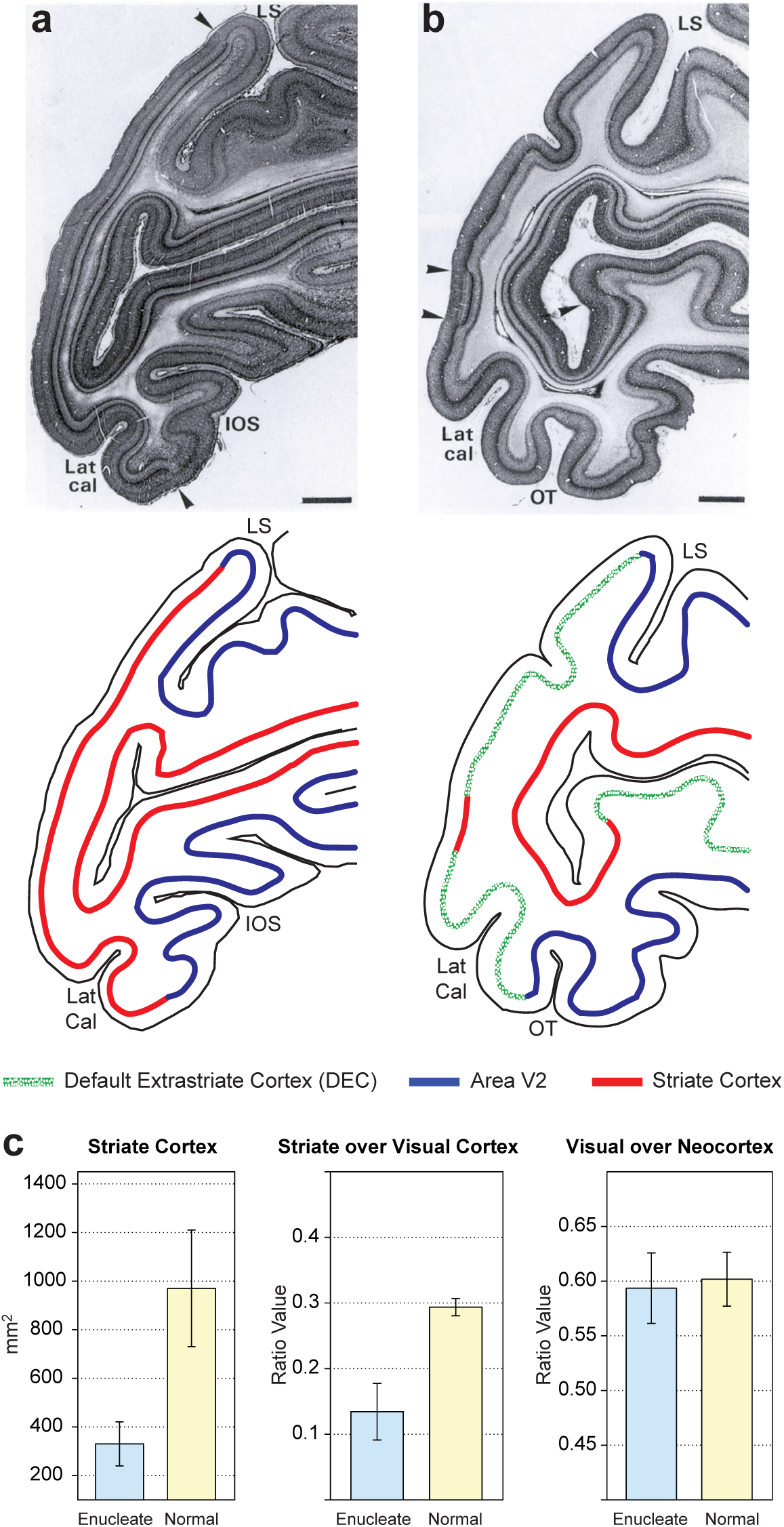
Effects of early enucleation on cortical parcellation. (**A**) Upper panel parasaggital Nissl stained sections showing cytoarchtecture; lower panel schematic showing the limits of striate cortex and area V2; (**B**) upper panel, parasaggital Nissl stained section in the neonate following prenatal enucleation at 68 days after conception (E68); lower panel, limits of areas V1, V2 and default extrastriate cortex. Sections in A and B taken from equivalent levels, arrow heads indicate limits of striate cortex. Note, large reduction of striate cortex on operculum and more modest reduction in the calcarine, scale bars, 2 mm. (**C**) Quantitative effects of enucleation on proportions of visual cortex; left-hand panel, surface area of striate cortex (p = 4.04e-05, 7 enucleates, 6 normals); middle-panel, proportion of striate cortex with respect to total visual cortex (p = 3.04e-06, 7 enucleates, 6 normals); right-hand panel, proportion of visual cortex with respect to total cortex (p = 0.63, 6 enucleates, 6 normals); Error bars, SD.

We used retrograde tracers that allow exploring the intrinsic labelling of a cortical area, which corresponds to the local connectivity(*15*). In the normal brain intrinsic connectivity represents 80-90% of the total connectivity and exhibits an exponential decline with distance(*15, 16*). In the anophthalmic brain the space constant of the exponential decline is significantly larger and intrinsic projections extend over considerable distances (**Fig. 2A**). Hence in the normal V1 the 75% threshold is at 0.85mm, the 80% at 0.95mm and the 95% at 1.7mm. In the DEC these distances are increased 4 to 6 fold (**Fig. 2B**), so that local connectivity extends across a large extent of the DEC on the operculum (**Fig. S1**).

**Fig. 2.**
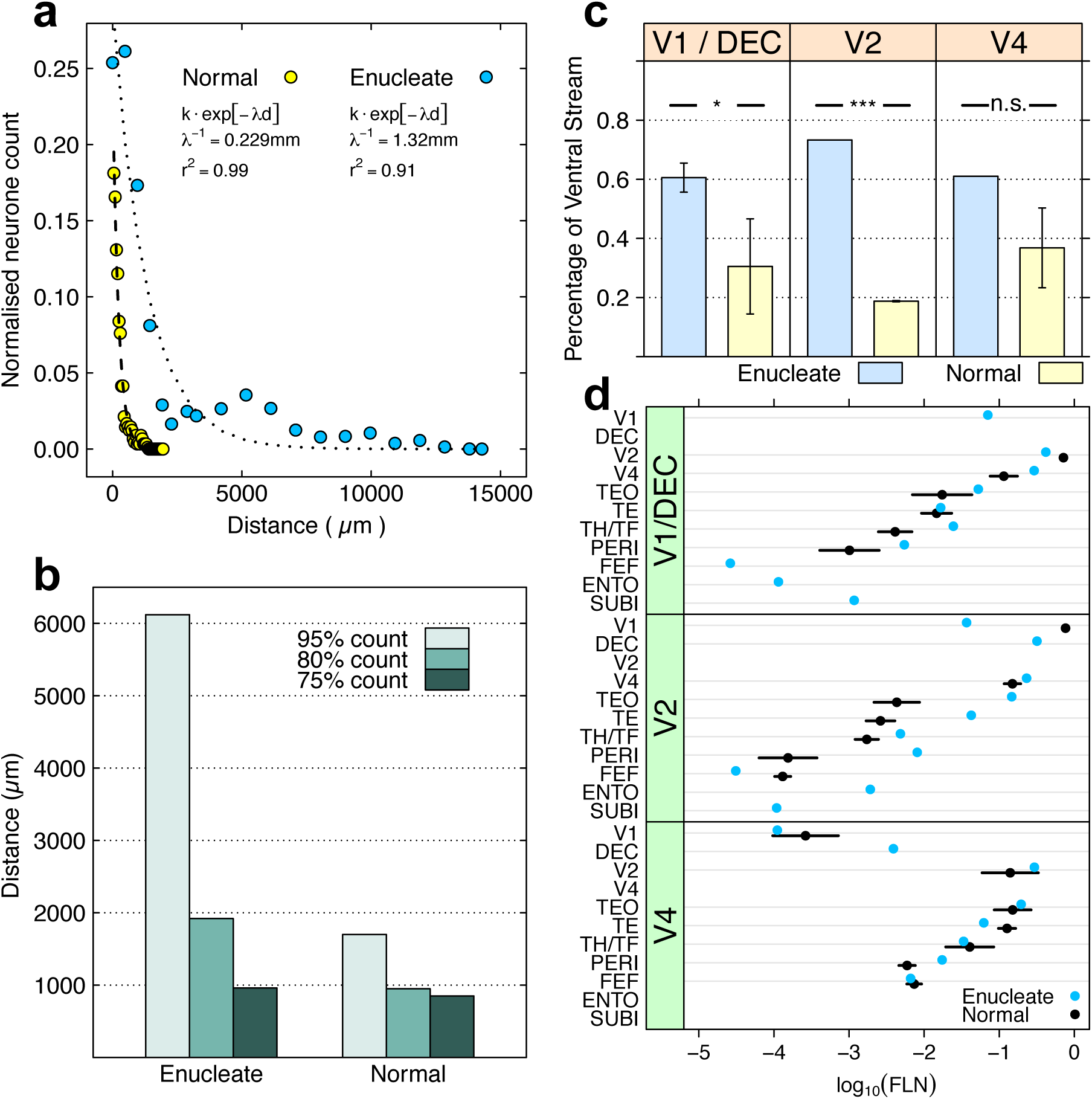
Intrinsic connectivity following enucleation and effects of ventralization. (A) Exponential decay of density of intrinsic neurons with distance following injection in area V1 of a normal (yellow dots, dashed line fit) and in the DEC (enucleation at E73) (blue dots, dotted line fit). The dashed and dotted lines represent exponential fits. **(B**) Distances within which the 3 thresholds (75%, 80%, and 95%) of intrinsic labelling are attained in normal V1 and in the DEC (enucleation at E73). **(C**) mean cumulative sum of Fraction of Labelled Neurons (FLN) in ventral stream areas; far-left panel, injections in area V1 and default extrastriate cortex (DEC; p = 0.0155); middle panel, injections in area V2 (p < 2e-16); right-hand panel, injections in area V4 (p = 0.301). Enucleate vs. normal across all injection: p = 2.52e-04; error bars, SD. All tests were performed assuming that the proportions followed a beta distribution(*39*). **(D**) Effect of enucleation on connexion strength in ventral stream areas. Log scale dot plot of FLN. Enucleates, blue dots; normal controls, black dots; upper-panel, injections in normal striate cortex (V1) and Default Extrastriate Cortex (DEC) (1 enucleate, 5 normals); middle-panel, injections in area V2 (1 enucleate, 3 normals); bottom panel, injections in area V4 (2 enucleates, 3 normal); error bars, SD. For abbreviations of area names see glossary.

The topography of connectivity in the anophthalmic was overall similar to that in the normal cortex. Following retrograde tracer injection in a target area, the numbers of labelled neurons in a given source area with respect to the total number of labelled neurons in the brain defines the Fraction of Labelled Neurons FLN, a weight index reflecting the strength of the particular pathway(*15*). High frequency sampling of labelled neurons in the cortex allows characterization with a single injection of the weighted connectivity of a pathway linking any two cortical areas(*17*). Injections limited to the grey matter were performed in DEC, V2 and V4 in three anophtalmic brains (**Fig. S2; Table S1**). Inspection of cortical labelling suggested that early visual deafferentation leads to an increase of numbers of labelled neurons in ventral stream areas (**Fig. S2, Fig S3**). Injections in DEC and area V2 show that deafferentation profoundly affects the relative strengths of the dorsal and ventral pathways, as seen after summing the FLN values across all ventral versus dorsal stream areas (**Fig. 2C, D**). Differences between normal and anophthalmic cumulative FLN values were not found to be significant following injections in area V4, suggesting that the effects of deafferentation are restricted to early cortical areas.The laminar distribution of retrogradely labelled parent neurones of a pathway is defined by its proportion of supragranular labelled neurons or SLN index, which has been shown to be highly consistent across individuals(*11*). The SLN values of a pathway define it as feedforward or feedback and specify a hierarchical distance(*18*). In the absence of the retina there is an increase in numbers of labelled supragranular layer neurons (**Fig. S4**). The SLN value for area V2 projections to DEC is significantly higher than for the V2 projection to V1 and likewise the projection of V3, FST and PIP to DEC have significantly increased SLN values (**Fig S4A**).Following deafferentation, projections to area V2 showed significant increases in the SLN in areas V1 and V3 as well as the dorsal stream areas MT, V3A, LIP, PIP, STP and PGa as well as an increase in the ventral stream area TEO (**Fig. S4B**). By contrast, deafferentation had little or no effect on the SLN values for any of the projections to area V4 (**Fig. S4D**) with the marked exception of V1 where only infragranular neurons were observed. However, given the very low FLN value, this cannot be considered significant. The most marked change in the SLN is the projection of area V1 to the DEC (**Fig. 3A**). Labelled neurons in area V1 projecting to the DEC are entirely located in the supragranular layers, which makes this projection very different from any projection from area V1 in the normal brain (**Fig. 3A**). Area V1 also projects strongly to area V2, but the V1->V2 pathway originates from both infra- and supragranular layers. A projection of V1 which is entirely from the supragranular layers would be to area V4, but the V1->V4 pathway is considerably weaker than the DEC->V4 pathway.

**Fig. 3.**
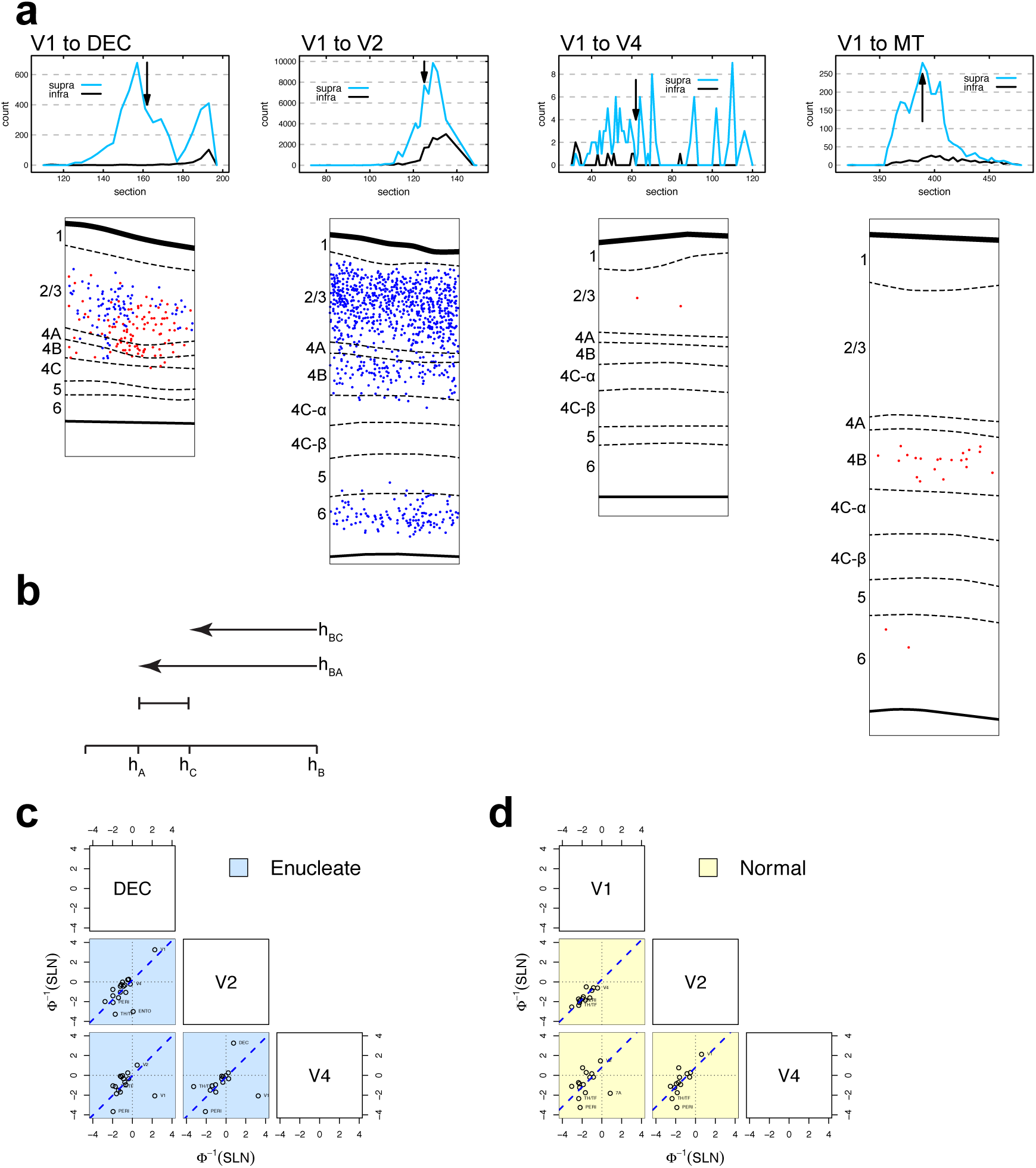
Laminar distribution of labeled cells and hierarchy. (A) High power plots comparing laminar distributions of V1->DEC to projections in the normal cortex. This comparison shows that the V1->DEC has no counter part in the normal cortex, either in terms of strength of projection nor laminar distribution. Black arrows indicate the section used for high power plots. **(B) and (C**) Pairs plots between probit-transformed SLN values of common areas from injections in V1/DEC, V2 and V4 (as indicated in the boxes along the diagonal). Each point represents the average pair of SLN values obtained in a single source area; the blue dashed lines indicate the best fitting lines of unit slope. **(B**) Enucleated cases (blue background). **(C**) Data from normal controls (yellow background). Area labels for points of potential interest and outliers are indicated to the right of the point.

SLN has been shown to be a robust indicator of hierarchical distance(*11, 19*) (see Materials and Methods). When the SLN values extracted from injections at different levels are mapped on to hierarchical space by means of a sigmoid function they display a surprising regularity, shown by the probit transformed values of SLN plotted in a pairwise fashion; if the transformation leads to a coherent measure of hierarchical distance the points will cluster around lines of unit slope (see Materials and Methods, **Fig. S6**). Importantly, both the normal and anophthalmic brains display this consistency in laminar organization (**Fig. 3C, D**). All the common projections to areas V1 and V2 in the normal cortex are feedback and, as expected, are observed in the lower left quadrant. This contrasts with the anophthalmic where V1 is in the top right quadrant, indicating it to be feedforward to both DEC and V2. Consideration of an ensemble of SLN values following injections in areas V1, V2 and V4 allows fitting a hierarchical model to both the normal and anophthalmic data sets (**Fig. 4A,** see Materials and Methods)(*11*). This shows that the overall layout of the ventral and dorsal stream areas remain approximately similar to that observed in the normal. However, in the anophthalmic brain, DEC and area V2 are considerably higher in the hierarchy than expected. The goodness of fit of the model is shown by close agreement between the empirical and model estimated SLN values by source and target areas (**Fig. 4B**, normal r^2^=0.72; enucleate r^2^=0.67).

**Fig. 4.**
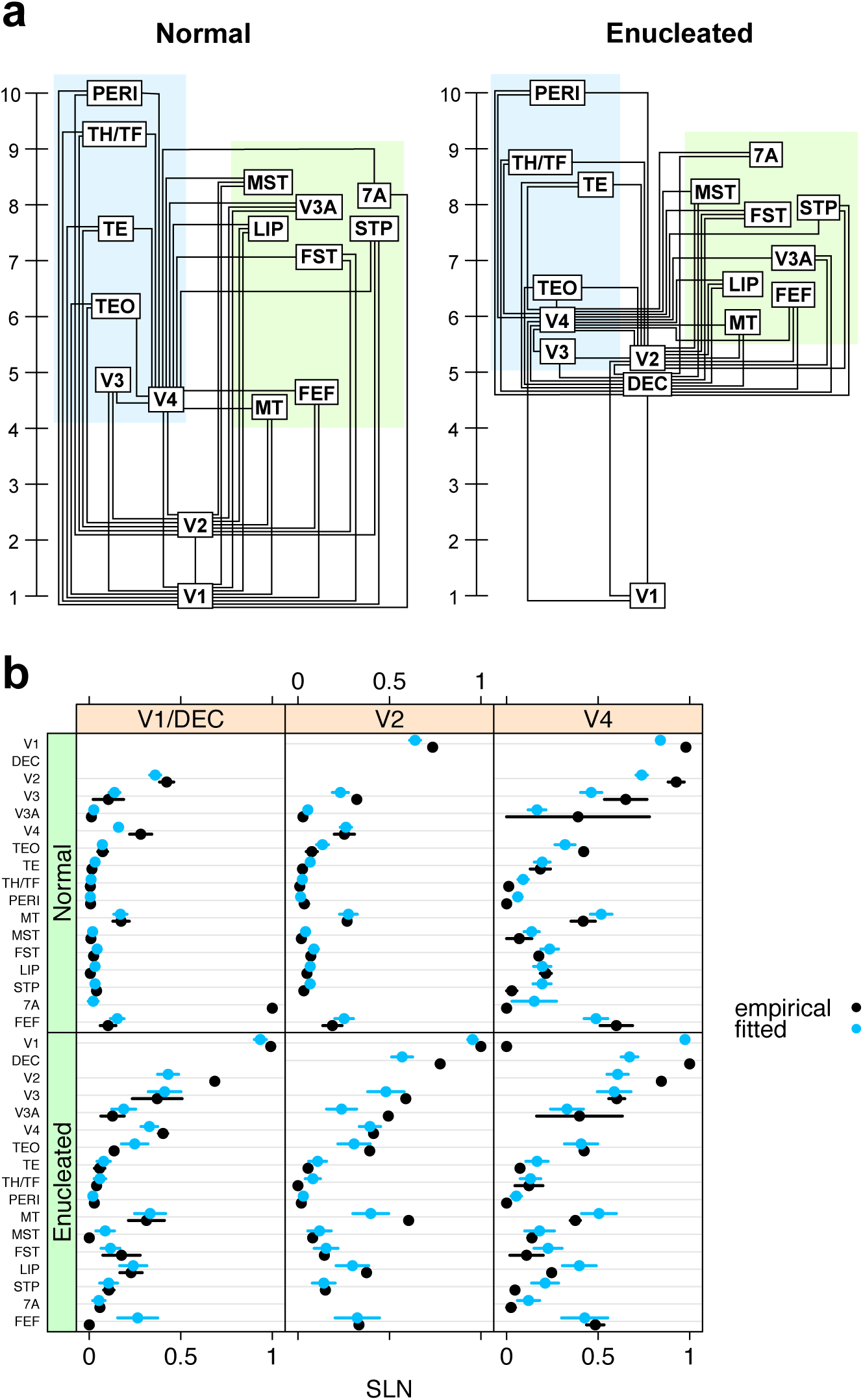
Developmental plasticity of the visual hierarchy. (A) Hierarchical relationship between areas in the visual system based on injections in V1, V2 and V4 in normal animals, and following enucleation and injections in Default Extrastriate Cortex (DEC), V2 and V4. **(B**) Detailed comparison of the model with the data for each injection by labelled area; black dots, empirical values; blue squares, predicted values. Error bars are empirical SEs for the data and model based SEs for the predictions.

The present results show that in the anophthalmic the topography of connectivity and global organization of the ventral and dorsal streams are largely conserved as has been suggested by imaging studies in the human congenitally blind(*20–26*), in line with the evidence of early developmental specification of the functional streams(*27, 28*). Further, in the anophthalmic brain we observe en expansion of the ventral pathway that could reflect cross modal plasticity(*24*).The present findings show that early primate visual cortex in anophthalmia is profoundly modified both in its cytoarchitecture and local connectivity. While the global hierarchy remains largely conserved, there are important local changes in the hierarchical organization. These changes would seem to reflect a persistence of immature features making the congenitally blind ‘visual’ cortex neotenic. Indeed, interareal connectivity during *in utero* development in the primate undergoes extensive remodelling characterized by a greatly expanded population of supragranular projecting neurons, a global hierarchical organization similar to that found in the adult, an absence of ectopic pathways and finally a markedly extensive local connectivity(*11, 29–32*). The relatively high position in the cortical hierarchy and the conservation of an extensive local connectivity in the DEC could ensure the long time constants which would be required for the observed higher cognitive functions of the deafferentated cortex of the blind(*19, 33*).The present findings need to be considered in view of current understanding of the developmental specification of the cortex. Developmental patterning of the neocortex is consequent to an interplay between intrinsic genetic mechanisms based on morphogens and secreted signalling molecules and extrinsic inputs relayed to the cortex by thalamocortical axons(*34–36*). The role of thalamic axons in arealization is a multistep hierarchical process involving events at progenitor and neuronal levels(*37*). A recent spatiotemporal transcriptome analysis of the pre- and postnatal macaque forebrain revealed a small number of genes that have persistent expression across cortical development, suggesting a large potential for extrinsic shaping of the cortex(*38*). Interestingly, this study shows that areal and laminar molecular phenotypes are acquired late postnatally indicating a wide and potentially important role of contextual shaping of the structure and function of the cortex, suggesting that particular attention should be paid to the care of the young congenitally blind.

## Supplementary Materials

Material and Methods

Figures S1-S7

Tables S1-S2

References (*39–47*)

